# Positive Affect Modulates Early Valuation and Conflict Processing in Social Decision-Making

**DOI:** 10.64898/2026.03.05.709732

**Authors:** Zhengxian Liu, Yunting Liu, Weixian Li, Cui Ruifang, Xiaobo Liu

**Author notes:** means corresponding author: E-mail: Ruifang Cui, Xiaobo Liu.

## Abstract

Social decision-making relies on dynamic affect–cognition interactions across distributed brain networks, yet how incidental positive affect modulates these mechanisms at a millisecond timescale remains unclear. This study investigated the impact of music-induced positive emotion on the neural dynamics of decision-making in the Ultimatum Game. Fifty-six participants were assigned to either a happy music group or an active control (rain sound) group. Fifty-six participants were assigned to either a happy music group or an active control (rain sound) group, while electroencephalography was recorded to capture rapid neural dynamics. Behaviorally, happy music accelerated reaction times (RTs) and decoupled the ERP–RT correlations observed in the control condition. Neurally, positive affect amplified event-related potential amplitudes during early conflict detection (220–280 ms) and late valuation (520–560 ms) stages. Multivariate pattern analysis further revealed that happy music enhanced the neural separability and temporal stability of decision states (accept vs. reject). Moreover, using support vector regression based on functional network features, we found that decision acceptance rates were predicted with significantly higher accuracy in the happy music group (*R* = 0.60) compared to controls (*R* = 0.41). Crucially, feature weight analysis indicated a topological shift in decision strategy: while the control group relied on frontal–central edges (implicating executive control), the happy music group was characterized by central–temporal connections (suggesting integrative processing). Collectively, these findings provide novel evidence that incidental emotion intervenes at the millisecond timescale to bias social choices, offering a dynamic network-based account of the affect-cognition interaction.

## Introduction

Social decision-making is a fundamental aspect of human interaction, requiring a dynamic interplay of cognitive evaluation and affective states, rather than a purely rational process of utility maximization (Rahal & Fiedler, 2022; Terenzi et al., 2021). The ultimatum game (UG) provides a well-established setting for studying this dynamic, in which individuals must weigh financial gain against social norms by accepting or rejecting a monetary offer (Sanfey et al., 2003). Neuroimaging studies have linked UG choices to a distributed conflict-control network including the anterior insula and lateral prefrontal cortex, reflecting affective appraisal and control processes engaged during offer (Rilling & Sanfey, 2011; Sanfey et al., 2003; Xu et al., 2025). However, decisions are rarely made in an emotional vacuum. Growing evidence suggests that incidental emotions—affective states unrelated to the decision itself—can bias these economic/social choices (Bartholomeyczik et al., 2022; Pertl et al., 2024). As a pervasive and potent inducer of affect, music offers a unique ecological window into how positive mood modulates social judgments and choice tendencies (Curzel et al., 2024; Nummenmaa et al., 2021; Sachs et al., 2021). Yet, the precise neural dynamics through which musical affect reconfigures the decision-making process remains under-characterize.

Theoretically, the mood maintenance hypothesis posited that positive affect motivated individuals to sustain their benevolent state, leading to increased cooperativeness and a higher propensity to accept offer (Isen, 2001). Complementary to this, the affect infusion model suggested that positive mood promoted a heuristic processing style, reducing the reliance on rigorous, bottom-up scrutiny of potential norm violations (Forgas, 1995; Harlé & Sanfey, 2007). Despite the robustness of these behavioral effects, the underlying neural mechanisms remain debated. Does positive affect shift the decision-making strategy toward a different neural pathway?

To unravel the temporal dynamics underlying this affective modulation, electroencephalography (EEG) offers millisecond-level resolution essential for tracking the rapid stages of decision formation (Hewig et al., 2011). Previous research has identified components such as the medial frontal negativity or feedback-related negativity (MFN/FRN) in the approximately 200–350 ms range and late positive potential (LPP) as markers of early conflict monitoring and late valuation, respectively (Mothes et al., 2016; Wu et al., 2011). However, traditional univariate event-related potential (ERP) analysis captures only a fraction of the information distributed across the scalp and often fails to reveal how the brain distinguishes between decision states (i.e., “Accept” vs. “Reject”) over time. Multivariate pattern analysis (MVPA) combined with machine learning classifiers can decode whole-brain ERP topographies at each time point, thereby revealing when distinct decision states begin to diverge (Kragel et al., 2018). Meanwhile, temporal generalization analysis can illuminate the temporal stability of these representations—determining whether the decision process relies on a transient, evolving code or a stable, sustained attractor state (Bode et al., 2011; Gold & Heekeren, 2014; King & Dehaene, 2014).

Furthermore, complex social decision-making relies on the integration of information across distributed networks (Bassett & Sporns, 2017). Thus, investigating how positive affect alters functional connectivity during the early stages of decision-making is crucial. Functional connectivity metrics, such as phase-locking value (PLV), capture spatial synchronization patterns that support different stages of decision-making (Lachaux et al., 1999). Ultimately, standard group-level statistics often overlook individual variability. Machine learning techniques, such as support vector regression (SVR) or other multivariate decoding methods applied to EEG time series, offer a powerful means to predict individual decision tendencies (e.g., acceptance or fairness evaluations) from neural patterns (Bode et al., 2022; Saeidi et al., 2021). It has been demonstrated by multivariate EEG decoding approaches that link trial-level neural signals to choice behavior in economic and social decision paradigms (Moerel et al., 2025; Moore et al., 2021).

Thus, by integrating ERPs, MVPA, functional connectivity, and machine learning techniques, the present study soughed to elucidate the neural dynamics through which positive musical affect modulated social decision-making in the UG. We hypothesize: (1) Happy music would speed up response times, consistent with evidence that positive musical stimulation improved processing speed and reduced reaction times (RTs) in cognitively demanding tasks by enhancing mood and arousal (Forgas, 2013; Orpella et al., 2025); (2) Positive affect would modulate both early conflict-monitoring (MFN) and late evaluative (LPP) ERP components, reflecting an enhanced sensitivity to social signals at multiple processing stages (Farkas & Sabatinelli, 2023; Mothes et al., 2016; Wu et al., 2011); (3) Using MVPA and temporal generalization analysis, happy music would enhance the separability and temporal stability of “Accept” vs. “Reject” neural patterns. This would suggest that positive affect sharpened the neural code or facilitates a more stable decision state (Takacs et al., 2020); and critically, (4) Positive affect would instantiate a distinct cognitive strategy that could be detected as a unique predictive network topology. Specifically, we predicted a transition from a frontal-executive control network (characteristic of neutral/calculative processing) to a central-temporal integration network (reflecting heuristic or semantic processing) in the happy condition (Geng et al., 2024; Gholam Tamimi & Daliri, 2025). By integrating these multidimensional features, this study aimed to provide a comprehensive mechanistic account of how happy music affect biases social economic choices.

### Materials and methods Data acquisition

A total of 56 healthy, right-handed undergraduate students were recruited from the University of Electronic Science and Technology of China. To control for potential confounds in auditory processing, individuals with formal musical education or extensive instrument training were excluded. Participants were randomly assigned to one of two experimental conditions (n=28 per group): (1) the happy music group (15 male, age 22.75±1.58), who listened to happy music; and (2) the active control group (15 male, age 22.82±2.48), who were exposed to rain sound stimulus.

The study protocol was approved by the Ethics Committee of the University of Electronic Science and Technology of China and conducted in strict accordance with the Declaration of Helsinki. Written informed consent was obtained from all participants prior to the experiment, and they were monetarily compensated for their participation.

### Procedure

In alignment with Juslin et al, the happy music group listened to a validated positive musical excerpt (No. HAPPY01) from the Chinese Affective Music Scale (CAMS) to induce positive emotions (Juslin et al., 2008). While the active control group listened to a recording of rain sounds, serving as a neutral auditory baseline. The experimental session lasted approximately one hour. Upon arrival, participants first completed a digital survey regarding their musical preferences and training background to ensure eligibility. They were then provided with detailed instructions for the entire experiment and completed a practice block to ensure task comprehension. Following preparation for EEG recording, a 5-minute eyes-closed resting-state baseline was acquired, followed by the formal UG task which lasted approximately 15 minutes.

To prevent dual-task interference and ensure that cognitive resources were not diverted from the decision-making task (Thompson et al., 2012), a sequential design was employed where the auditory induction preceded the behavioral task. Specifically, participants were exposed to the stimulus (music or rain) for one minute. To verify the efficacy of the mood induction, participants completed a brief emotional assessment before (T1) and after (T2) the listening phase using a 5-point Likert scale, such as “Currently, I am in a good mood,” “As I answer these questions, I feel cheerful,” “For some reason, I am not very comfortable right now,” (reverse-scored) and “At this moment, I feel sad” (reverse-scored) (Underwood & Froming, 1980). The cumulative score represented an emotion score, with higher values indicating elevated emotional well-being. Cronbach’s alpha indicated acceptable internal consistency for the mood scores at both time points (T1: α=0.668; α=0.665). A paired sample *t*-test confirmed a significant increase in positive affect scores in the happy music group following the exposure (*p* < 0.001).

Upon completing the questionnaires, a gender-matched confederate entered the laboratory. The participant and confederate were seated at separate tables to prevent visual contact but allowed for auditory interaction. The confederate was then escorted to a separate room under the pretense of acting as the “Proposer,” while the participant was assigned the role of “Responder.” In reality, all offers were computer-generated and predetermined. As illustrated in Figure 1, each trial began with a fixation cross (“+”) presented for 800 ms, followed by a display of the total stake (¥10) for 500 ms. After a 1000 ms black screen interval, one of the five predetermined allocation options was presented: fair (¥5:¥5), moderately unfair (¥6:¥4; ¥7:¥3), and extremely unfair (¥8:¥2; ¥9:¥1). Each options appearing randomly 40 times for a total of 200 trials. Participants accepted or rejected the offer by pressing the “F” or “J” keys (counterbalanced across participants) using their index fingers. Stimulus presentation and response recording were executed using E-Prime 3.0 software (Psychology Software Tools, Pittsburgh, PA).

**Figure 1.**
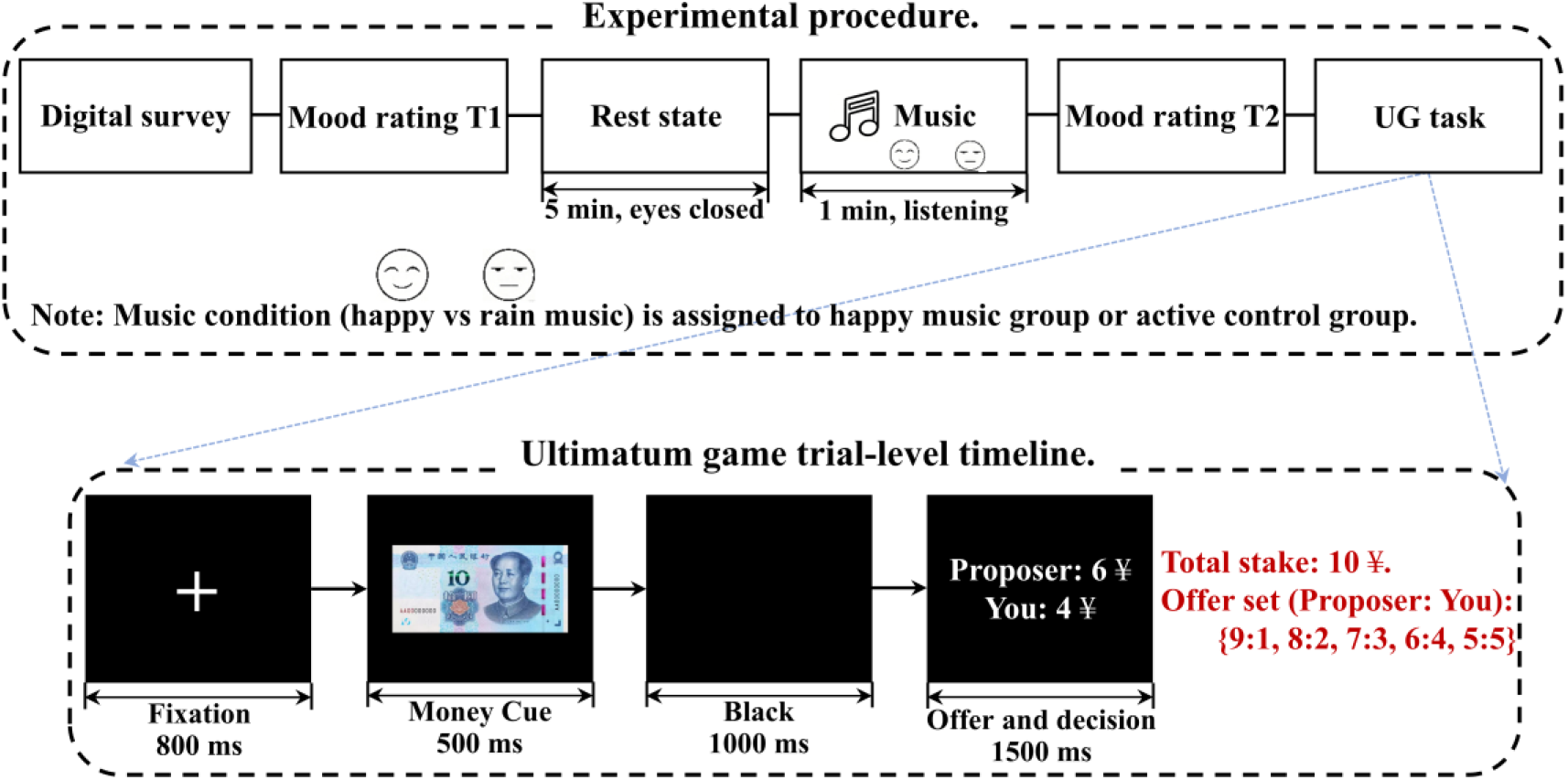
The experimental design.

### EEG Data recording

EEG signals were recorded using 64-channel Ag/AgCl electrode cap (ANT Neuro, Germany) arranged according to the extended 10-20 system. Signals were digitized at a sampling rate of 1000Hz with an online bandpass filter of 0.3–100Hz, using CPz as the online reference. Electrooculograms (EOGs) for monitoring eye movements were concurrently recorded from an extra channel situated on the left eye. Electrode impedances were kept below 10 kΩ throughout the session.

### Data preprocessing

Offline data analysis was performed using the Neuroscience information toolbox (NIT) (Dong et al., 2018). First, raw EEG data underwent visual inspection to identified identify malfunctioning channels, which were substituted with extrapolated average values derived from neighboring 3–4 channels (Quan et al., 2023). Subsequently, The continuous data were then segmented into resting-state and task-state blocks. For each state, the EEG data were transformed into average reference data and subjected to filtering with a passband of 0.5–20 Hz, along with a notch filter (45–55 Hz). Re-referencing of the EEG data to the “zero” reference was then performed using the reference electrode standardization technique (REST) offline (Yao, 2001). Artifacts arising from ocular movements (blinks, saccades) and muscle activity were identified and removed, ensuring that only artifact-free data were retained for subsequent analyses.

### Event-Related-Potential and Behavior Analysis

The clear EEG was segmented into epochs ranging from -200 ms to 800 ms that 0 ms was the offer onset. Epochs were then baseline-corrected using the -200 to 0 ms pre-stimulus interval. An automatic artifact rejection procedure was applied, excluding any trials where the signal amplitude exceeded ±60 μV. Fz, FC1, FCz, FC2, and Cz electrodes were collapsed and averaged as an indication of frontal activity based on prior literature and inspection of the grand average waveforms (Kappenman & Luck, 2010). Finally, cluster-based permutation tests (cluster-forming threshold *p* < 0.05) were conducted to comparing differences between two music groups.

Behavioral data analysis primarily examined group differences in reaction times (RTs) among three music conditions using paired-sample *t* tests. Furthermore, to explore the neuro-behavioral link, Pearson correlation analyses were conducted between the amplitudes of ERP components showing significant group differences and the corresponding decision-making reaction times.

### Single-trial multivariate pattern analysis (MVPA)

To evaluate the impact of music on the neural dynamics of decision-making, multivariate pattern classification was employed using linear support vector machines (SVM) via the *fitcsvm* and *predict* functions in MATLAB. The classifier was trained to discriminate between decision outcomes (“Accept” vs. “Reject”) based on the spatial topography of the EEG signal at each time point. These analyses were conducted separately for the happy music and active control groups. We hypothesized that if happy music promotes with the cognitive processing required for decision-making, this would be reflected in an increase of decoding accuracy (i.e., improved distinctiveness of neural patterns) compared to the control condition. For each participant, the SVM classification was performed at each time point along the epoch, utilizing amplitude data from all EEG channels as features. To ensure the robustness of the results, a stratified cross-validation procedure was implemented. The training and testing processes were iterated 100 times, and the resulting classification accuracies were averaged to produce a final temporal decoding course.

To further elucidate the temporal organization of information-processing stages, a Temporal Generalization Analysis was conducted (Dehaene & King, 2016). Unlike standard time-resolved decoding, this method assesses whether neural representations are transient or sustained over time. Specifically, a SVM classifier trained on data from a specific time point (t) was tested on data from all other time points (t^′^). A 10-fold cross-validation scheme was utilized, and the procedure was repeated 100 times to compute a generalization matrix for each subject. Then we evaluated performance of classifier via accuracy.

To determine whether classification performance significantly exceeded the chance level (50%), one-sample *t*-tests were computed across the time series. Family-wise errors (FWE) were controlled using cluster-based permutation tests. A null distribution was generated by randomly shuffling the classification labels 1,000 times. Clusters were initially identified using a threshold of *p* < 0.05. The observed cluster masses (sum of *t*-statistics) were then contrasted against the surrogate distribution derived from the permutation procedure. Only clusters falling within the 95th percentile (significance threshold *p* < 0.05, corrected) were considered statistically significant. Finally, to allow for the neurophysiological interpretation of the decision-making states, the weight vectors of the linear classifiers were transformed into activation patterns using the method described by Haufe et al. (Haufe et al., 2014).

### Network analysis

Network based on the early stage of decision making (0 – 300 ms) was constructed via phase locking value (PLV) (Aydore et al., 2013), which could measure the phase synchronization between two narrow-band signals. PLV between signals *s1(t)* and *s2(t)* was formulized as follows.

First, the Hilbert transform was used:

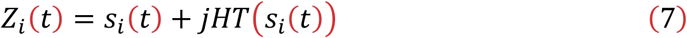

Where *HT*(*si*(*t*)) is the Hilbert transform of *si(t*)defined as

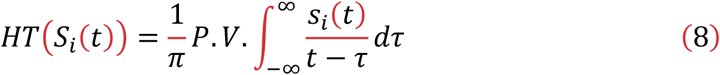

And P.V. denotes Cauchy principal value. Once the analytic signals are defined, the constant phase between *Z*_1_(*t*) and *Z*_2_(*t*) can be computed as

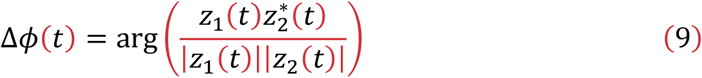

The instantaneous PLV is then defined as (Lachaux, et al., 1999; Celka, 2007)

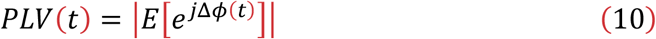

Where *E[*.] represents the expected value. Nine regions of interest were selected and the difference map based on PLV of different groups was calculated.

Finally, to validate the functional significance of the initial decision-making stage, an SVR analysis was implemented to predict the behavioral acceptance rate (Liu et al., 2021; Pan et al., 2021; Si et al., 2020). The SVR models were trained using functional network features derived from the averaged ERP trials. A 10-fold cross-validation scheme was applied separately for the happy music and active control groups, with the entire training and testing procedure iterated 100 times to ensure stability. The predictive performance of the regression models was quantified by comparing the predicted values against the actual acceptance rates using four metrics: pearson’s correlation coefficient (*R*), the coefficient of determination (*R*^*2*^), root mean squared error (*RMSE*), and mean absolute error (*MAE*) (Liu, Huang, et al., 2025; Liu, Xu, et al., 2025). Furthermore, to identify the specific neural connections driving the prediction, the relative importance of each network edge was evaluated using a Random Forest algorithm (Liu et al., 2021).

## Results

### Experimental Paradigm and Mood Induction

To validate the efficacy of the mood manipulation protocol, self-reported mood ratings were collected using a 5-point Likert scale at two distinct time points: immediately before (T1) and after (T2) the one-minute listening task (happy music or white noise). Paired-sample *t*-tests were conducted to assess changes in emotional state. A significant increase in positive affect was observed following exposure to happy music (*p* < 0.001), whereas no significant changes were detected in the white-noise condition. Consequently, the experimental manipulation was confirmed to have successfully modulated positive affect, ensuring the validity of the subsequent neuro-behavioral analyses.

### ERP Amplitude Differences

ERPs were extracted from the frontal lobe included Fz, FC1, FCz, FC2, and Cz electrodes within a time windows of −200 to 800 ms time-locked to the onset of the offer. As shown in Figure 2, cluster-based permutation testing (cluster-forming threshold *p* < 0.05, *p*_*peak*_=3.85 × 10^-21^, *t* =9.45) revealed that, compared with the active control group, the happy music group exhibited significantly larger ERP amplitudes during both an early window (220–280 ms) and a late window (520–560 ms).

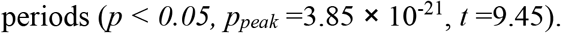

**Figure 2.**
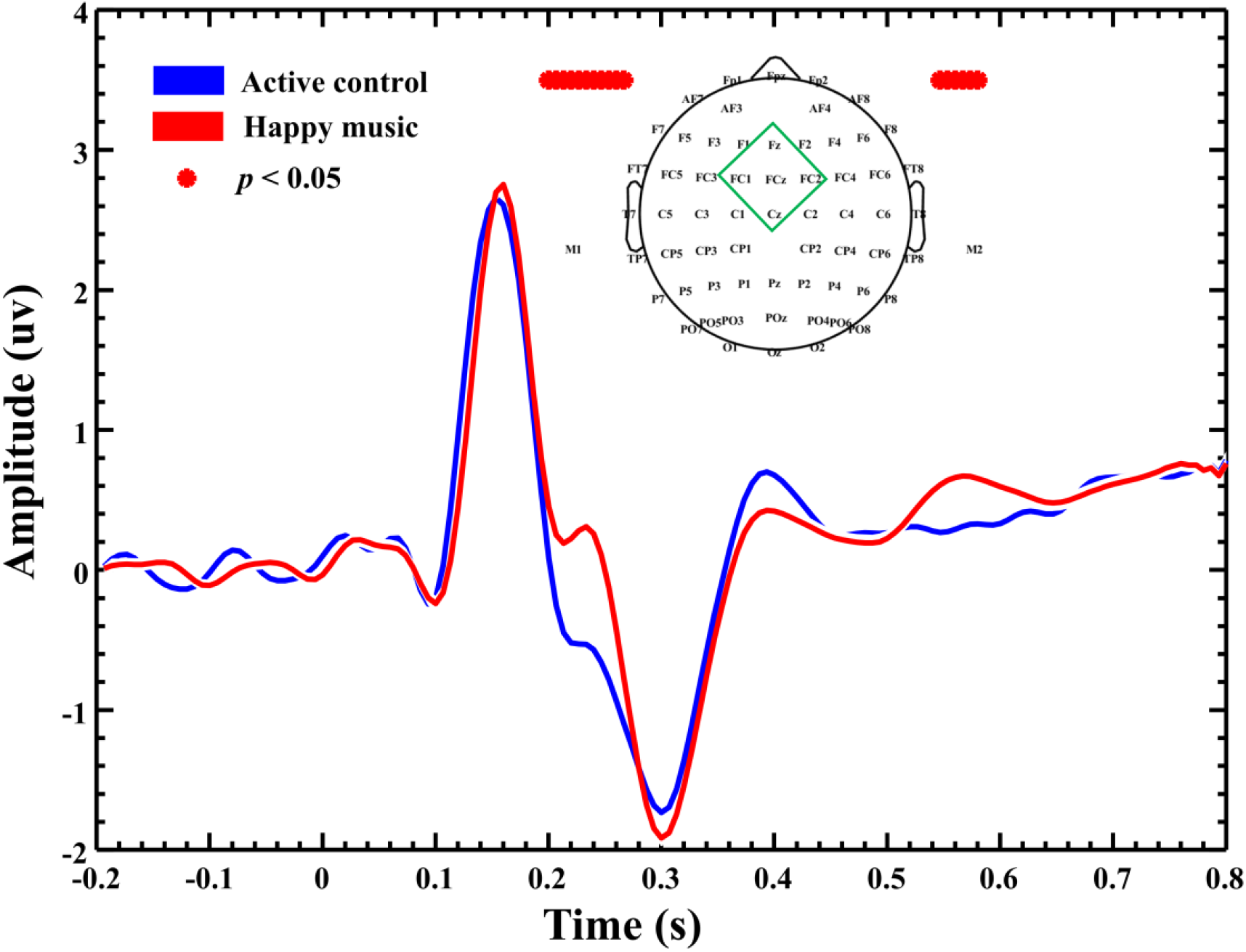
ERP waveforms corresponding to the decision acceptance condition were derived the frontal lobe included Fz, FC1, FCz, FC2, and Cz electrodes. Distinct differences in amplitude were observed primarily during the early and late latency

### Behavioral Reaction Times and ERP–RT Correlation

RTs, averaged across 200 trials per participant, were analyzed using independent-samples *t*-tests. Significantly shorter response time were observed in the happy music group compared to the active control group (*p* < 0.05, see Figure 3A). These results suggest that the speed of social decision-making is facilitated by the induction of positive affect.

**Figure 3.**
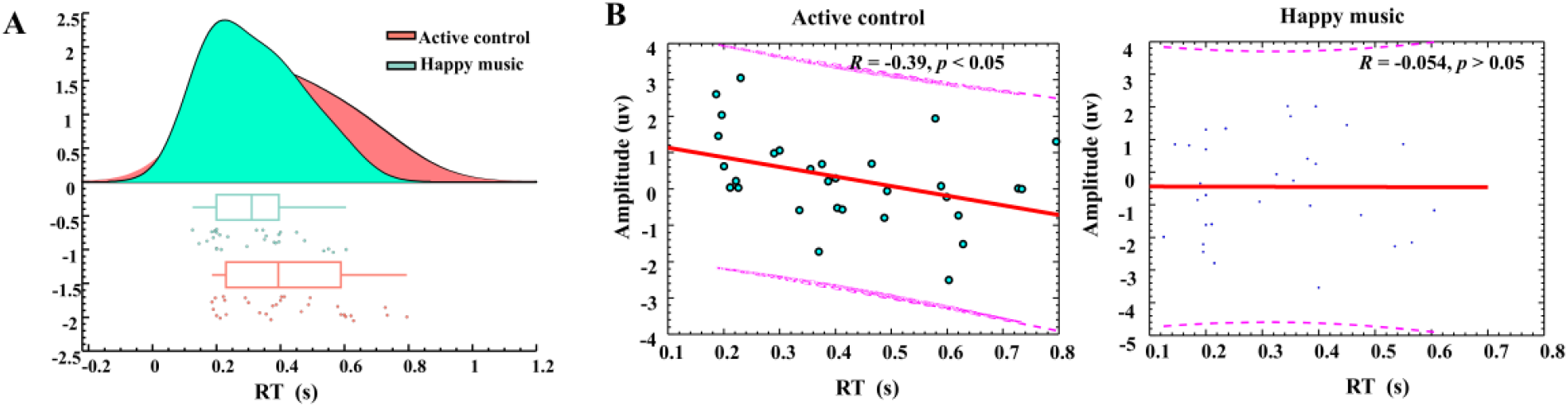
Analysis of behavioral performance and neuro-behavioral correlations. A. Comparison of reaction times (RTs) between the happy music and active control groups. Error bars represent standard error of the mean (SEM). B. Scatter plots illustrating the correlation between mean ERP amplitudes and RTs for both the active control (*R = -0*.*39, p < 0*.*05*) and happy music groups (*R = 0*.*054, p > 0*.*05*).

As shown in Figure 3B, regarding the relationship between neural activity and behavior, a significant negative correlation was identified within the active control group between mean ERP amplitude in the early time window (220–280 ms) and RTs (*R* = –0.39, *p* = 0.031), which larger early neural responses were found to be associated with faster decision speeds. Conversely, no statistically significant relationship between ERP amplitudes and RTs was detected in the happy music group (*p* > 0.05). This dissociation implies that the direct coupling between neural signal magnitude and decision latency may be modulated or decoupled by the presence of positive affect.

### Temporal MVPA Decoding Accuracy

As illustrated in Figure 4, a linear SVM classifier was trained at each time point to distinguish between “Accept” and “Reject” trials using single-trial ERP spatial topographies. This procedure employed 10-fold cross-validation with 100 repetitions. Above-chance decoding accuracy (> 50%) was achieved by both groups within the 200–300 ms latency range. Furthermore, subsequent cluster-based statistical analysis demonstrated that decoding accuracy in the happy music group was significantly higher than that of the active control group within the 230 to 440 ms time window (*p* < 0.05). These results indicate that the discriminability of neural patterns underlying social decision outcomes is enhanced by the induction of positive affect.

**Figure 4.**
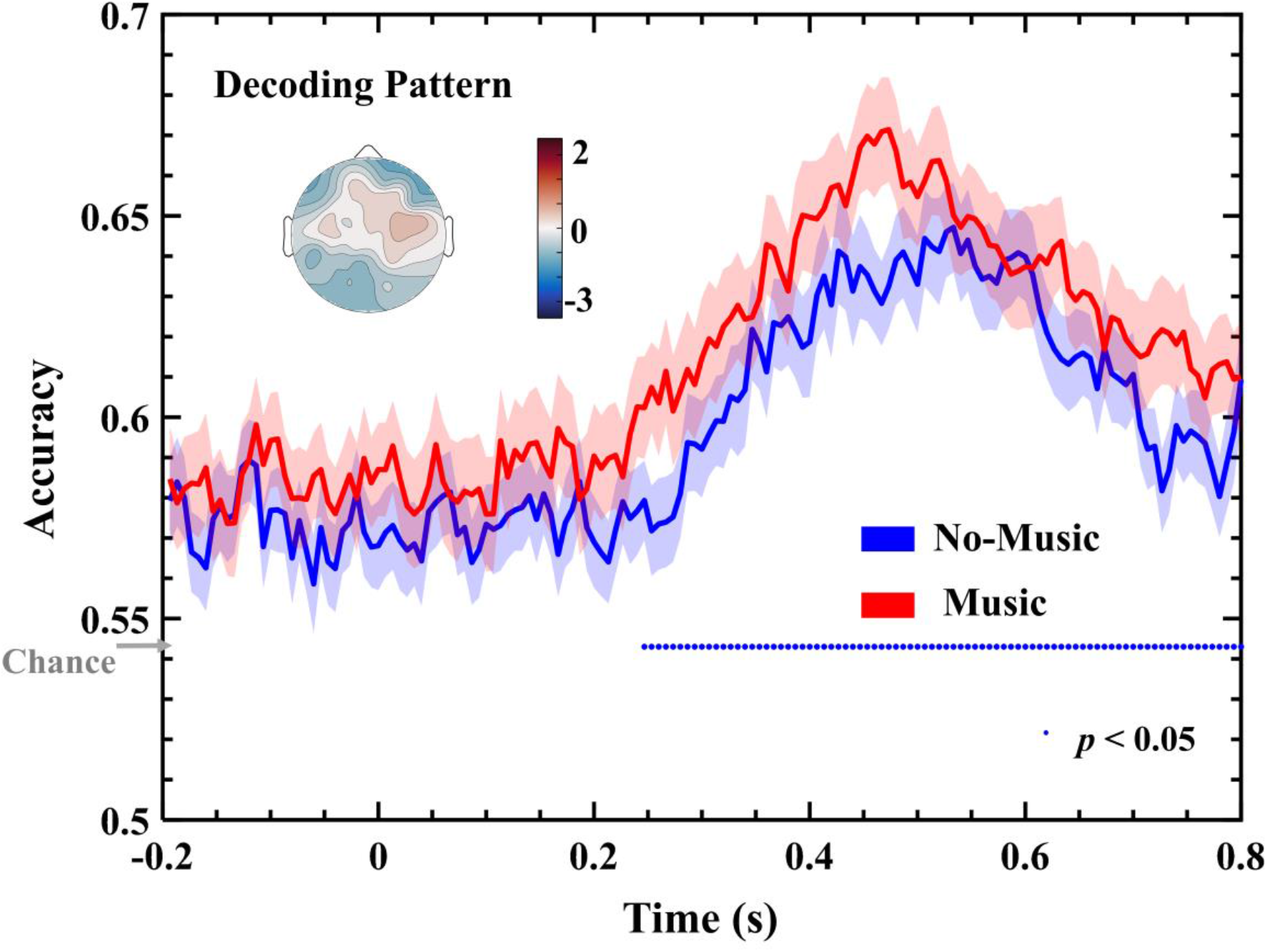
Linear SVM classification analysis to decode “Accept” and “Reject” trials. A 10-fold cross-validation procedure was implemented to decode decision outcomes (Accept vs. Reject) at each time point using ERP spatial topography features. The blue dots indicate time points where decoding accuracy is significantly above the chance level (50%).

Temporal generalization matrices were computed to examine how classifiers trained at a specific time point generalize to other time points, as shown in Figure 5. A diagonal band of above-chance accuracy was observed in both groups, indicating the presence of temporally sustained neural representations. Furthermore, a significant cluster between 230 and 440 ms (T > cluster threshold) was identified in the difference matrix (music minus active control). This finding confirms that not only is instantaneous decoding boosted by the musical stimulus, but representational stability across time is also enhanced.

**Figure 5.**
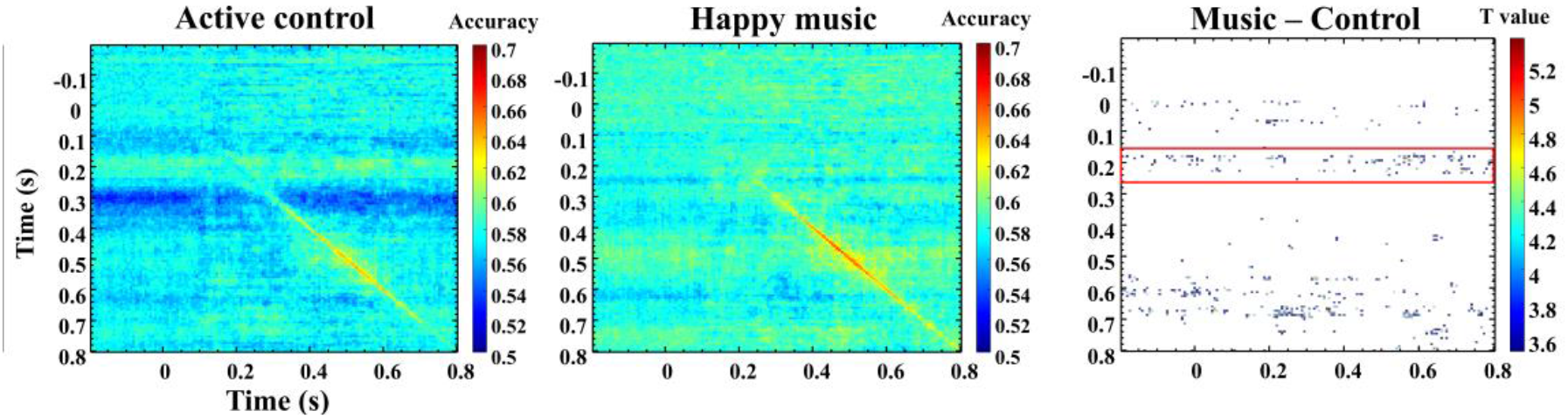
Temporal generalization of decoding accuracy. Temporal generalization matrices displaying decoding performance for the happy music and active control groups. The axes represent the training time (y-axis) and testing time (x-axis). The difference matrix highlights time windows where representational stability significantly differed between the two groups (*p*_*fdr*_ *< 0*.*05*).

### Prediction of Acceptance Rates from Early-Stage Functional Networks

As depicted in Figure 6, functional networks were constructed based on PLVs between electrode pairs during the early decision-making window (0–300 ms), which were utilized as input features for an SVR model to predict the acceptance rate of each participant. A predictive correlation of *R* = 0.60 (*R*^*2*^ = 0.35), *MAE* = 0.09, *RMSE* = 0.11, was achieved in the happy music group (*p* < 0.05). Meanwhile, a correlation of *R* = 0.41 (*R*^*2*^ = 0.14), *MAE* = 0.14, *RMSE* = 0.16 was observed in the active control group (*p* < 0.05).

**Figure 6.**
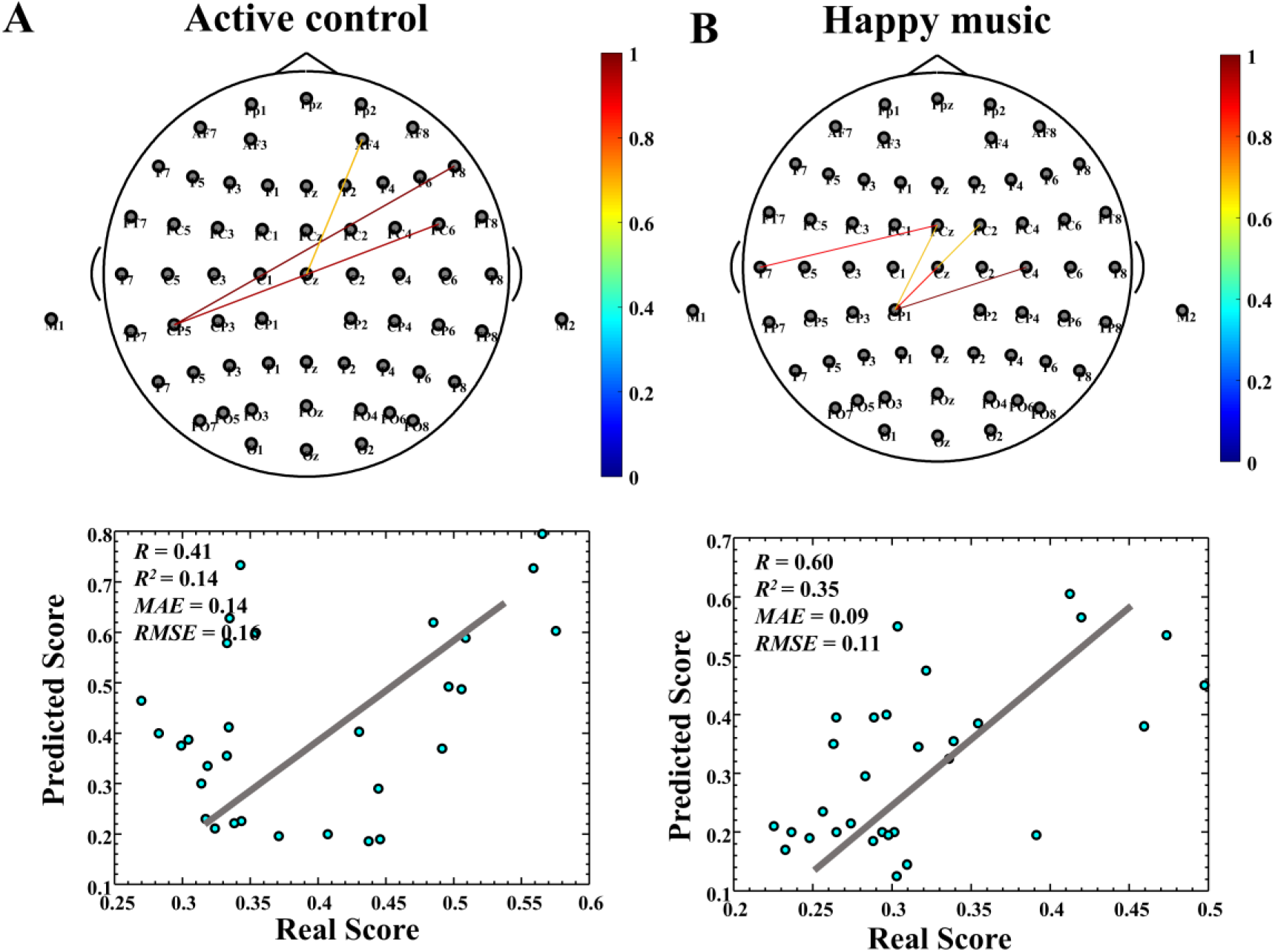
SVR-based behavioral prediction using network features. Prediction of acceptance rates using functional network features derived from the initial stage of decision-making in the happy music (*R* = 0.60, *MAE* =0.09, *p* < 0.05) (A) and active control groups (*R* = 0.41, *MAE* =0.14, *p* < 0.05) (B).

Through the topographical visualization of feature weights, it was found that predictions in the happy music group relied primarily on central–temporal connections, whereas frontal–central edges were identified as the key features in the active control group. These results demonstrate that social decision preferences are not only reflected by early-stage network dynamics but can also be robustly predicted by them, particularly under the influence of positive affect.

## Discussion

The present study investigated the neuro-computational mechanisms by which incidental positive affect reconfigures social decision-making. Behaviorally, happy music expedited response times, indicating a facilitation of the decision process. Neurophysiologically, happy music biases decision dynamics by strengthening the early neural encoding of value and conflict, which accompanied by amplified MFN and late positivity. Beyond these univariate changes, MVPA revealed that happy music increased the separability and temporal stability of neural states distinguishing “accept” from “reject” decisions. Crucially, SVR analysis uncovered a fundamental shift in the predictive network topology: whereas decisions in the control group relied on frontal-central connectivity, those in the happy group were robustly predicted by central-temporal connectivity. Collectively, these findings suggest that positive affect intervenes at the very outset of the decision process, tuning the neural dynamics to alter how offers are evaluated and acted upon.

Behaviorally, participants under music responded more quickly, consistent with theories that positive mood accelerates cognitive flexibility and conflict resolution (Isen, 2001). The shortened RTs in the happy group further suggest a shift toward a more fluent, heuristic processing mode, consistent with the notion that positive mood reduces the cognitive load typically associated with deliberative bargaining (Forgas, 2013). In addition, the larger midline frontal negativities under happy music aligns with literature linking the MFN/FRN component to fairness conflict detection in social exchanges (Hewig et al., 2011; Wu et al., 2011). The augmentation of early ERP amplitude suggests that positive affect potentiates the neural salience of unfair offers, perhaps by introducing a parallel reward signal that “buffers” negative conflict processing. The subsequent increase in late-stage amplitude likely reflects deeper reappraisal or emotion-regulation operations (Boksem & De Cremer, 2010), indicating that music not only amplifies immediate conflict signals but also modulates the downstream evaluative cascade. These finding suggested that positive affect may heighten neural gain, enhancing the signal-to-noise ratio of incoming social stimuli (Eldar et al., 2016). Crucially, correlational analyses revealed a dissociation: while the control group showed a significant negative correlation between early ERP amplitude and RTs (larger neural response predicted faster decisions), this relationship was absent in the happy group. In the control condition, the decision process appears to follow a sequential accumulation model, where the strength of early conflict detection directly drives the speed of the motor output (Gluth et al., 2020). Conversely, the decoupling observed in the happy group suggests a shift toward a more heuristic or automatic processing mode (Forgas, 1995). Under positive affect, the decision to accept may be driven by a generalized pro-social prior, making the behavioral output less dependent on the trial-by-trial magnitude of early neural conflict monitoring. In addition, this decoupling parallels findings in emotional Stroop paradigms where mood induction disrupts direct ERP–behavior mappings (Schirmer & Kotz, 2006), supporting the notion that music creates an alternative neural trajectory for transforming conflict signals into action.

Multivariate pattern analysis confirmed that both groups exhibit above-chance separability of accept versus reject states in the 200–300 ms window, yet music extended this window of high decoding accuracy and improved cross-temporal generalization. This result aligns with the broaden-and-build theory at a neural level, suggesting that positive emotion expands the distinctiveness of cognitive repertoires (Fredrickson, 2001). Furthermore, the sustained diagonal pattern in temporal generalization under happy music suggests that positive affect not only sharpens momentary neural distinctions but also promotes more durable representational formats. Stable neural representation implies that once the brain enters a decision state, it maintains that configuration robustly, rather than rapidly fluctuating between states (King & Dehaene, 2014). This attractor state stability likely facilitates the faster reaction times observed: the happy brain ‘locks in’ to the decision outcome more efficiently, reducing the neural noise that typically prolongs deliberation in neutral or ambiguous contexts.

By leveraging PLV to capture early functional connectivity, our SVR model predicted acceptance rates more robustly under music, highlighting central-temporal links as critical predictors. This finding extends network-based decision-prediction work (Park & Friston, 2013) by showing that an affective manipulation selectively amplifies certain edges. The more significant finding was the dissociation in predictive networks. In the active control group, acceptance rates were predicted by frontal-central functional connectivity. This topography maps onto the canonical “cognitive control” network, involving the medial frontal cortex and lateral prefrontal regions. These areas are extensively documented in resolving the conflict between self-interest and fairness norms (Cutler & Campbell-Meiklejohn, 2019). The reliance on this network implies that under neutral music conditions, the decision to accept is a result of “calculative control”—actively overriding the impulse to reject. In contrast, the happy music group exhibited a predictive reliance on central-temporal connections. The temporal lobes, particularly the superior temporal sulcus and temporal pole, are critical hubs for social cognition, supporting mentalizing and the integration of social-semantic information (Olson et al., 2013; Schurz et al., 2021). These findings suggests that positive music may reconfigure the computational basis of the decision-making. Instead of a conflict resolution process (frontal regions), the brain may adopt an integrative strategy (temporal regions), where the offer is processed through pathways associated with social meaning and emotional coherence. This network switch provides a neural explanation for the affect infusion model (Forgas, 1995), indicating that positive mood allows social information to be processed more integratively and less critically.

The above results demonstrated that these network features may be established rapidly (0–300 ms post-stimulus) and possess high predictive power (*R* = 0.60 for the happy music group). It suggests that affective context primes specific functional networks early in the processing stream, constraining the subsequent behavioral output. The higher prediction accuracy in the happy group compared to controls further implies that positive affect may induce a more stable, low-noise neural state, making behavior more consistent and predictable from brain activity.

Several limitations warrant consideration. First, only a single “happy” music stimulus was employed, limiting the ability to generalize across different emotional tones or musical genres; future work should compare multiple affective categories and cultural styles. Second, the sample comprised healthy young adults performing the UG, so the findings require replication in more diverse populations and decision paradigms to establish broader applicability. Third, while EEG offers excellent temporal resolution, its spatial precision is limited; combining EEG with fMRI or advanced source-localization techniques would help localize the underlying neural circuitry. Fourth, music was presented only prior to the task, leaving open questions about the effects of continuous or task-embedded auditory interventions. Finally, although we identified robust neurobehavioral associations, causal mechanisms remain to be tested, for instance through brain-stimulation methods targeting conflict-monitoring networks.

## Conclusion

In summary, the present study demonstrates that positive affect fundamentally reconfigures the neuro-computational architecture of social decision-making. The results showed that happy music does not merely amplify early neural signatures of conflict detection and valuation, but crucially enhances the separability and temporal stability of the underlying decision states. Most significantly, the decision states were underpinned by a topological shift in predictive networks: happiness prompts a transition from a frontal-executive control mode to a temporal-integrative mode. This reorganization suggests that positive affect facilitates social cooperation by engaging efficient, heuristic neural pathways rather than resource-heavy cognitive control. Collectively, these findings provide novel evidence that incidental emotion intervenes at the millisecond timescale to bias social choices, offering a dynamic network-based account of the affect-cognition interaction.

## Acknowledgement

This work is supported by .

## References

Aydore, S., Pantazis, D., & Leahy, R. M. (2013). A note on the phase locking value and its properties. Neuroimage, 74, 231–244.

Bartholomeyczik, K., Gusenbauer, M., & Treffers, T. (2022). The influence of incidental emotions on decision-making under risk and uncertainty: a systematic review and meta-analysis of experimental evidence. Cognition and Emotion, 36(6), 1054–1073.

Bassett, D. S., & Sporns, O. (2017). Network neuroscience. Nature Neuroscience, 20(3), 353–364. 10.1038/nn.4502

Bode, S., He, A. H., Soon, C. S., Trampel, R., Turner, R., & Haynes, J.-D. (2011). Tracking the unconscious generation of free decisions using uitra-high field fMRI. PloS one, 6(6), e21612.

Bode, S., Schubert, E., Hogendoorn, H., & Feuerriegel, D. (2022). Decoding continuous variables from event-related potential (ERP) data with linear support vector regression using the Decision Decoding Toolbox (DDTBOX). Frontiers in Neuroscience, 16, 989589.

Boksem, M. A., & De Cremer, D. (2010). Fairness concerns predict medial frontal negativity amplitude in ultimatum bargaining. Social neuroscience, 5(1), 118–128.

Curzel, F., Osiurak, F., Trân, E., Tillmann, B., Ripollés, P., & Ferreri, L. (2024). Enhancing musical pleasure through shared musical experience. Iscience, 27(6), 109964.

Cutler, J., & Campbell-Meiklejohn, D. (2019). A comparative fMRI meta-analysis of altruistic and strategic decisions to give. NeuroImage, 184, 227–241.

Dehaene, S., & King, J.-R. (2016). Decoding the dynamics of conscious perception: the temporal generalization method. Micro-, meso-and macro-dynamics of the brain, 85–97.

Dong, L., Luo, C., Liu, X., Jiang, S., Li, F., Feng, H., Li, J., Gong, D., & Yao, D. (2018). Neuroscience information toolbox: An open source toolbox for EEG–fMRI multimodal fusion analysis. Frontiers in Neuroinformatics, 12, 56.

Eldar, E., Niv, Y., & Cohen, J. D. (2016). Do you see the forest or the tree? Neural gain and breadth versus focus in perceptual processing. Psychological Science, 27(12), 1632–1643.

Farkas, A. H., & Sabatinelli, D. (2023). Emotional perception: Divergence of early and late event-related potential modulation. Journal of Cognitive Neuroscience, 35(6), 941–956.

Forgas, J. P. (1995). Mood and judgment: the affect infusion model (AIM). Psychological Bulletin, 117(1), 39–66.

Forgas, J. P. (2013). Don’t worry, be sad! On the cognitive, motivational, and interpersonal benefits of negative mood. Current Directions in Psychological Science, 22(3), 225–232.

Fredrickson, B. L. (2001). The role of positive emotions in positive psychology: The broaden-and-build theory of positive emotions. American Psychologist, 56(3), 218–226.

Geng, H., Xu, P., Aleman, A., Qin, S., & Luo, Y.-J. (2024). Dynamic organization of large-scale functional brain networks supports interactions between emotion and executive control. Neuroscience Bulletin, 40(7), 981–991.

Gholam Tamimi, M., & Daliri, M. R. (2025). State-base dynamic functional connectivity analysis of fMRI data during facial emotional processing. Brain Imaging and Behavior, 19, 1307–1318.

Gluth, S., Kern, N., Kortmann, M., & Vitali, C. L. (2020). Value-based attention but not divisive normalization influences decisions with multiple alternatives. Nature Human Behaviour, 4(6), 634–645.

Gold, J. I., & Heekeren, H. R. (2014). Neural mechanisms for perceptual decision making. In Neuroeconomics (pp. 355–372). Elsevier.

Harlé, K. M., & Sanfey, A. G. (2007). Incidental sadness biases social economic decisions in the Ultimatum Game. Emotion, 7(4), 876–881.

Haufe, S., Meinecke, F., Görgen, K., Dähne, S., Haynes, J.-D., Blankertz, B., & Bießmann, F. (2014). On the interpretation of weight vectors of linear models in multivariate neuroimaging. Neuroimage, 87, 96–110.

Hewig, J., Kretschmer, N., Trippe, R. H., Hecht, H., Coles, M. G., Holroyd, C. B., & Miltner, W. H. (2011). Why humans deviate from rational choice. Psychophysiology, 48(4), 507–514.

Isen, A. M. (2001). An influence of positive affect on decision making in complex situations: Theoretical issues with practical implications. Journal of Consumer Psychology, 11(2), 75–85.

Juslin, P. N., Liljeström, S., Västfjäll, D., Barradas, G., & Silva, A. (2008). An experience sampling study of emotional reactions to music: listener, music, and situation. Emotion, 8(5), 668.

Kappenman, E. S., & Luck, S. J. (2010). The effects of electrode impedance on data quality and statistical significance in ERP recordings. Psychophysiology, 47(5), 888–904.

King, J.-R., & Dehaene, S. (2014). Characterizing the dynamics of mental representations: the temporal generalization method. Trends in Cognitive Sciences, 18(4), 203–210.

Kragel, P. A., Koban, L., Barrett, L. F., & Wager, T. D. (2018). Representation, pattern information, and brain signatures: from neurons to neuroimaging. Neuron, 99(2), 257–273.

Lachaux, J. P., Rodriguez, E., Martinerie, J., & Varela, F. J. (1999). Measuring phase synchrony in brain signals. Human brain mapping, 8(4), 194–208.

Liu, X., Huang, Y., Wang, Y., Lin, C., Huang, Y., & Liu, X. (2025). Self-supervised electrocardiograph de-noising. IEEE Access.

Liu, X., Xu, Y., Xu, H., He, L., Long, S., Huang, Y., Wang, Y., Lu, Y., Huang, Y., & Wu, J. (2025). Advancing interpretable cardiac disease diagnosis via a transformer-convolutional hybrid network on electrocardiograms. Engineering Applications of Artificial Intelligence, 152, 110675.

Liu, X., Yang, S., & Liu, Z. (2021). Predicting Fluid Intelligence via Naturalistic Functional Connectivity Using Weighted Ensemble Model and Network Analysis. NeuroSci, 2(4), 427–442.

Moerel, D., Grootswagers, T., Chin, J. L., Ciardo, F., Nijhuis, P., Quek, G. L., Smit, S., & Varlet, M. (2025). Neural decoding of competitive decision-making in Rock–Paper–Scissors. Social Cognitive and Affective Neuroscience, 20(1), Nsaf101.

Moore, M., Katsumi, Y., Dolcos, S., & Dolcos, F. (2021). Electrophysiological correlates of social decision-making: an eeg investigation of a modified ultimatum game. Journal of Cognitive Neuroscience, 34(1), 54–78.

Mothes, H., Enge, S., & Strobel, A. (2016). The interplay between feedback-related negativity and individual differences in altruistic punishment: An EEG study. Cognitive, Affective, & Behavioral Neuroscience, 16(2), 276–288.

Nummenmaa, L., Putkinen, V., & Sams, M. (2021). Social pleasures of music. Current Opinion in Behavioral Sciences, 39, 196–202.

Olson, I. R., McCoy, D., Klobusicky, E., & Ross, L. A. (2013). Social cognition and the anterior temporal lobes: a review and theoretical framework. Social Cognitive and Affective Neuroscience, 8(2), 123–133.

Orpella, J., Bowling, D. L., Tomaino, C., & Ripollés, P. (2025). Effects of music advertised to support focus on mood and processing speed. PloS One, 20(2), e0316047.

Pan, H., Liu, X., Cai, X., & Lai, Y. (2021). Classification of schizophrenia EEG based on gamma-band brain network. International Journal of Psychophysiology, 168, S130–S131.

Park, H.-J., & Friston, K. (2013). Structural and functional brain networks: from connections to cognition. Science, 342(6158), 1238411.

Pertl, S. M., Srirangarajan, T., & Urminsky, O. (2024). A multinational analysis of how emotions relate to economic decisions regarding time or risk. Nature Human Behaviour, 8(11), 2139–2155.

Quan, L., Liu, X., Cui, R., Li, X., Liu, C., Yang, R., Gong, D., & Dong, L. (2023). Interaction effects of Action Real-Time S trategy Game experience and trait anxiety on brain functions measured via EEG rhythm. Brain-Apparatus Communication: A Journal of Bacomics, 2(1), 2176004.

Rahal, R.-M., & Fiedler, S. (2022). Cognitive and affective processes of prosociality. Current Opinion in Psychology, 44, 309–314.

Rilling, J. K., & Sanfey, A. G. (2011). The neuroscience of social decision-making. Annual Review of Psychology, 62(1), 23–48.

Sachs, M. E., FeldmanHall, O., & Tamir, D. I. (2021). Clarifying the link between music and social bonding by measuring prosociality in context. Behavioral and Brain Sciences, 44, e90.

Saeidi, M., Karwowski, W., Farahani, F. V., Fiok, K., Taiar, R., Hancock, P. A., & Al-Juaid, A. (2021). Neural decoding of EEG signals with machine learning: a systematic review. Brain Sciences, 11(11), 1525.

Sanfey, A. G., Rilling, J. K., Aronson, J. A., Nystrom, L. E., & Cohen, J. D. (2003). The neural basis of economic decision-making in the ultimatum game. Science, 300(5626), 1755–1758.

Schirmer, A., & Kotz, S. A. (2006). Beyond the right hemisphere: brain mechanisms mediating vocal emotional processing. Trends in cognitive sciences, 10(1), 24–30.

Schurz, M., Radua, J., Tholen, M. G., Maliske, L., Margulies, D. S., Mars, R. B., Sallet, J., & Kanske, P. (2021). Toward a hierarchical model of social cognition: A neuroimaging meta-analysis and integrative review of empathy and theory of mind. Psychological Bulletin, 147(3), 293–327.

Si, Y., Li, F., Duan, K., Tao, Q., Li, C., Cao, Z., Zhang, Y., Biswal, B., Li, P., & Yao, D. (2020). Predicting individual decision-making responses based on single-trial EEG. NeuroImage, 206, 116333.

Takacs, A., Mückschel, M., Roessner, V., & Beste, C. (2020). Decoding stimulus–response representations and their stability using EEG-based multivariate pattern analysis. Cerebral Cortex Communications, 1(1), tgaa016.

Terenzi, D., Liu, L., Bellucci, G., & Park, S. Q. (2021). Determinants and modulators of human social decisions. Neuroscience & Biobehavioral Reviews, 128, 383–393.

Thompson, W. F., Schellenberg, E. G., & Letnic, A. K. (2012). Fast and loud background music disrupts reading comprehension. Psychology of Music, 40(6), 700–708.

Underwood, B., & Froming, W. J. (1980). The mood survey: A personality measure of happy and sad moods. Journal of personality Assessment, 44(4), 404–414.

Wu, Y., Leliveld, M. C., & Zhou, X. (2011). Social distance modulates recipient’s fairness consideration in the dictator game: An ERP study. Biological psychology, 88(2-3), 253–262.

Xu, T., Zhang, L., Zhou, F., Fu, K., Gan, X., Chen, Z., Zhang, R., Lan, C., Wang, L., & Kendrick, K. M. (2025). Distinct neural computations scale the violation of expected reward and emotion in social transgressions. Communications Biology, 8(1), 106.

Yao, D. (2001). A method to standardize a reference of scalp EEG recordings to a point at infinity. Physiological measurement, 22(4), 693.

